# Human CCDC51 and yeast Mdm33 are functionally conserved mitochondrial inner membrane proteins that demarcate a subset of organelle fission events

**DOI:** 10.1101/2024.03.21.586162

**Authors:** Alia R. Edington, Olivia M. Connor, Madeleine Marlar-Pavey, Jonathan R. Friedman

## Abstract

Mitochondria are highly dynamic double membrane-bound organelles that exist in a semi-continuous network. Mitochondrial morphology arises from the complex interplay of numerous processes, including opposing fission and fusion dynamics and the formation of highly organized cristae invaginations of the inner membrane. While extensive work has examined the mechanisms of mitochondrial fission, it remains unclear how fission is coordinated across two membrane bilayers and how mitochondrial inner membrane organization is coupled with mitochondrial fission dynamics. Previously, the yeast protein Mdm33 was implicated in facilitating fission by coordinating with inner membrane homeostasis pathways. However, Mdm33 is not conserved outside fungal species and its precise mechanistic role remains unclear. Here, we use a bioinformatic approach to identify a putative structural ortholog of Mdm33 in humans, CCDC51 (also called MITOK). We find that the mitochondrial phenotypes associated with altered CCDC51 levels implicate the protein in mitochondrial fission dynamics. Further, using timelapse microscopy, we spatially and temporally resolve Mdm33 and CCDC51 to a subset of mitochondrial fission events. Finally, we show that CCDC51 can partially rescue yeast Δ*mdm33* cells, indicating the proteins are functionally analogous. Our data reveal that Mdm33/CCDC51 are conserved mediators of mitochondrial morphology and suggest the proteins play a crucial role in maintaining normal mitochondrial dynamics and organelle homeostasis.

## Introduction

Mitochondria are multifunctional double membrane-bound organelles whose shape and internal architecture are governed in response to cellular metabolic cues, nutrient availability, and respiratory demands (Bennett et al., 2022; Wai and Langer, 2016). While numerous processes give rise to mitochondrial network architecture, the overall distribution of the organelle arises from dynamic events, including cytoskeletal-based motility and inter-connectivity attributed to the ability of mitochondria to undergo opposing homotypic fusion and fission (Quintana-Cabrera and Scorrano, 2023; Ul Fatima and Ananthanarayanan, 2023). Additionally, the mitochondrial inner membrane (IMM) is dynamic and invaginates to form cristae that are home to membrane-shaping and ATP-producing oxidative phosphorylation machinery (Klecker and Westermann, 2021). Proper IMM architecture is required for optimal energy production, and the density of cristae membranes dynamically increases in response to increased respiratory demand (Balsa et al., 2019; Latorre-Muro et al., 2021). Thus, the metabolic functions of the organelle are intricately linked to the coordination of multiple dynamic shaping factors, processes, and machineries.

Despite our extensive knowledge of the mechanisms underlying mitochondrial dynamics and the identification of determinants of internal mitochondrial organization, we still have a poor understanding of how each is coordinated to maintain organellar homeostasis. In the case of mitochondrial fission, a conserved dynamin superfamily member (Drp1 in humans, Dnm1 in yeast) is recruited to the outer mitochondrial membrane (OMM), oligomerizes to encircle the organelle, and utilizes GTP hydrolysis to constrict and divide the membranes (Kraus et al., 2021). There is precise spatial control of mitochondrial fission, and most sites are pre-marked at the OMM by inter-organelle contacts with the endoplasmic reticulum (ER) (Friedman et al., 2011; Kleele et al., 2021). Mitochondrial fission is also spatially coordinated with mtDNA replication events that occur in the mitochondrial matrix (Lewis et al., 2016). Thus, fission must be coordinated across both the OMM and IMM, which raises the question of whether internal mitochondrial proteins are involved in or required for mitochondrial fission.

Recently, we identified the first internal mitochondrial protein that is required for mitochondrial fission in fungal species, the intermembrane space localized protein Mdi1 (also called Atg44) (Connor et al., 2023; Fukuda et al., 2023). Without Mdi1, Dnm1 can constrict the organelle but fails to complete fission of the OMM or IMM (Connor et al., 2023). As Mdi1 is not found in metazoan species, it remains an open question whether a functionally equivalent protein is required for division in higher eukaryotes. However, evidence that Dnm1 requires Mdi1 in yeast suggest that, in humans, Drp1 may not be able to sever both the IMM and OMM without the coordinated activity of internal factors. Another internal mitochondrial protein that was previously implicated in mitochondrial fission is the yeast protein, Mdm33 (also called She9) (Klecker et al., 2015; Messerschmitt et al., 2003). Mdm33 is an integral IMM protein that is required for normal mitochondrial morphology, and Δ*mdm33* yeast cells form elongated, hollow mitochondrial structures that are partially resistant to the induction of mitochondrial fission (Klecker et al., 2015; Messerschmitt et al., 2003). Additionally, Mdm33 has extensive coiled-coil domains on both the matrix and intermembrane space sides of the IMM, has been shown to self-associate and form high molecular weight structures, and is thus hypothesized to coordinate fission across the IMM (Messerschmitt et al., 2003). *MDM33* genetically interacts with other phospholipid synthesis coding-genes (Hoppins et al., 2011; Klecker et al., 2015). Thus, one proposed model is that Mdm33 coordinates mitochondrial lipid homeostasis pathways to influence mitochondrial architecture and/or fission dynamics (Klecker et al., 2015). However, Mdm33 is not thought to be conserved outside of fungal species, and its precise role remains elusive despite the unique phenotypes associated with its loss.

To begin to determine how mitochondrial architecture and dynamics are coordinated across the OMM and IMM in metazoans, we attempt to identify potential human functional analogs of Mdm33. Using a bioinformatic approach, we identify a structurally similar IMM protein, CCDC51. We find that loss of CCDC51 phenocopies mitochondrial defects of Δ*mdm33* yeast cells, and as in yeast, CDCC51 overexpression causes Drp1-dependent mitochondrial fragmentation. Interestingly, we observe that acute CCDC51 depletion leads to phenotypes associated with defects in mitochondrial fission, suggesting an involvement in the process. Using live cell microscopy, we find that both CCDC51 and Mdm33 are spatially and temporally linked to a subset of mitochondrial fission events. Finally, we show that CCDC51 can functionally complement yeast Δ*mdm33* cells, suggesting the proteins are indeed functionally orthologous. Thus, we have identified a human IMM protein that is required for the maintenance of mitochondrial morphology and may help facilitate mitochondrial dynamics in a conserved manner.

## Results

### Mitochondrial morphology defects in human CCDC51-depleted cells phenocopy yeast Mdm33 mutants

Despite the unique and severe mitochondrial morphology defects in yeast cells in the absence of Mdm33, the protein has no known ortholog in metazoans based on primary sequence homology. Mdm33 consists of an N-terminal mitochondrial targeting sequence (MTS) and two transmembrane domain segments that are interspersed with predicted coiled-coil domains (Messerschmitt et al., 2003) (Fig. 1A). Given that these Mdm33-defining domains are elements that may be under less evolutionary pressure to maintain primary sequence homology or may play a structural role (Surkont and Pereira-Leal, 2015; Truebestein and Leonard, 2016), we reasoned that a potential functional ortholog may exist with minimal sequence similarity. While a BLAST search failed to identify any metazoan sequence homologs of Mdm33, analysis using HHPRED (Zimmermann et al., 2018), which searches for remote homology, identified the human IMM protein CCDC51 as the top hit (E-value 4.9E-10; see Methods). Notably, CCDC51, also called MITOK based on its suggested role as a mitochondrial K^+^ channel, has been shown to be required for normal mitochondrial morphology (Paggio et al., 2019). CCDC51 is similar in length to Mdm33 (455 and 411 amino acids, respectively) and, like Mdm33, contains an N-terminal MTS and two transmembrane domain segments that are likewise interspersed with predicted coiled-coil domains (Fig. 1A).

**Figure 1.**
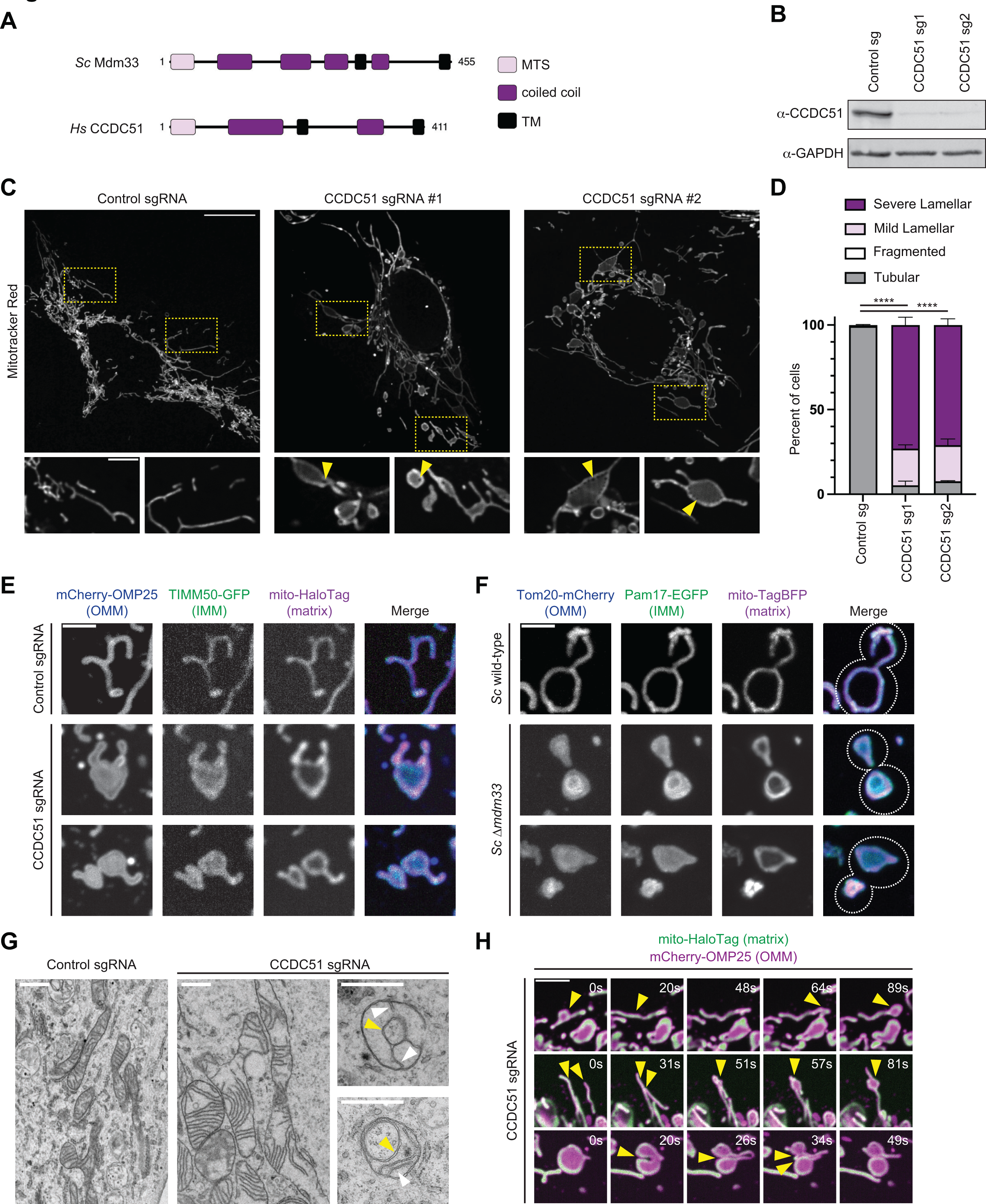
Mitochondrial morphology defects in human CCDC51-depleted cells phenocopy yeast Mdm33 mutants. (A) A schematic depicting the domain architecture of yeast Mdm33 and human CCDC51. **(B)** Western analysis with the indicated antibodies of whole cell lysates from U2OS CRISPRi expressing control sgRNA or sgRNAs targeting CCDC51. **(C)** Representative maximum intensity projection confocal images of U2OS CRISPRi cells expressing control or CCDC51-targeted sgRNAs and stained with Mitotracker Red. Insets below are single plane images and correspond to the indicated dashed boxes. Arrows mark discontinuities in Mitotracker staining. **(D)** A graph of mitochondrial morphology categorization from the indicated CRISPRi cells as in (C). Data shown represent 100 cells per condition in each of three independent experiments, and bars indicate S.E.M. Asterisks (****p<0.0001) represent unpaired two-tailed *t* tests of tubular mitochondrial morphology. **(E)** Representative maximum intensity projection confocal images are shown of the indicated CRISPRi cells expressing mCherry-OMP25, TIMM50-GFP, and mito- HaloTag labeled with JF646. **(F)** Maximum intensity projection confocal images are shown of the indicated yeast strains expressing Tom20-mCherry, Pam17-EGFP, and mito-TagBFP. Dashed lines indicate cell outlines. **(G)** Electron microscopy images are shown of mitochondria from U2OS CRISPRi cells expressing control or CCDC51 sgRNA, where indicated. Yellow arrows mark internal ring structures and white arrows mark cristae. **(H)** Single plane timelapse confocal microscopy images at the indicated intervals (s = seconds) are shown of U2OS CCDC51 CRISPRi cells expressing mCherry-OMP25 and mito-HaloTag/JF646. Arrows mark sites of resolution and reformation of lamellar mitochondria (top row), formation of lamellar mitochondria via apparent mitochondrial fusion (middle row), and fission of a lamellar mitochondria (bottom row). See also Movies S1-S3. Scale bars = (C)15 µm (5 µm in insets); (E, F) 3 µm; (G) 800 nm; (H) 5 µm.

The mitochondrial morphology defects of Δ*mdm33* cells are a defining characteristic, and to our knowledge, have not been observed in any other yeast mutant strains. Specifically, in the absence of Mdm33, mitochondria form extensive rings and sheet-like lamellar structures (Klecker et al., 2015; Messerschmitt et al., 2003). To explore the mitochondrial morphology of CCDC51- depleted cells in more detail and determine whether it is similar to that of Δ*mdm33* yeast cells, we utilized CRISPR interference (CRISPRi) to transcriptionally repress and stably deplete endogenous CCDC51 from U2OS cells, a human osteosarcoma cell line. U2OS cells stably expressing the dCas9-KRAB transcriptional repressor (Le Vasseur et al., 2021) were transduced with a lentiviral plasmid expressing scrambled sgRNA or individual sgRNAs targeting the transcription start site of CCDC51. We generated two different stable knockdown lines, each with near complete depletion of CCDC51 protein levels (Fig. 1B). We then examined mitochondrial morphology by confocal microscopy of cells stained with the dye, Mitotracker Red. In each CCDC51-depleted cell line, the mitochondria appeared to form lamellar, sheet-like structures, in contrast to the tubular morphology observed in control cells (Fig. 1C-1D). The Mitotracker labeling was also more intense at the edge of the lamellar mitochondrial structures, suggesting that the dye does not uniformly stain the mitochondria in these cells (Fig. 1C, inset). Additionally, we could occasionally observe small discontinuities in the mitochondrial staining that appeared as holes at the perimeter of lamellar structures (Fig. 1C, arrows). Thus, consistent with observations in HeLa CCDC51 knockout (KO) cells (Paggio et al., 2019), CCDC51 is required for normal mitochondrial morphology.

To further characterize the mitochondrial morphology defect of CCDC51-depleted cells, we transiently transfected cells expressing control sgRNA or CCDC51-targeted sgRNA with markers for the OMM (mCherry-OMP25), IMM (TIMM50-GFP), and matrix (mito-HaloTag labeled with JF646). In control cells, each fluorescent protein relatively uniformly labeled the mitochondrial membrane (Fig. 1E, top panel). In contrast, in CCDC51-depleted cells, each compartment marker had a unique localization pattern along the dilated lamellar structures. The matrix marker appeared similar to the Mitotracker staining, with more intense labeling at the periphery of the mitochondria and dimmer signal remaining in the interior (Fig. 1E, bottom panels). Interestingly, the matrix also appeared to form lariat ring structures, similar to previous observations in budding yeast Δ*mdm33* cells (Messerschmitt et al., 2003). Conversely, the IMM and OMM markers labelled lamellar mitochondrial structures more uniformly in a sheet-like manner (Fig. 1E, bottom panels). These markers appeared to encapsulate the non-uniform matrix marker, suggesting the appearance of the lariat-shaped mitochondria with either matrix markers or Mitotracker does not reflect the external structure of the organelle.

We then wanted to directly compare the mitochondrial morphology of mammalian CCDC51-depleted cells to yeast Δ*mdm33* cells. We chromosomally tagged yeast with functional markers of the OMM (Tom20-mCherry) and IMM (Pam17-EGFP) (Connor et al., 2023), and co-expressed the matrix marker, mito-TagBFP, in both wild-type and Δ*mdm33* cells. As expected, mitochondrial morphology was altered in Δ*mdm33* cells and the matrix marker formed hollow- appearing rings in many cells (Fig. 1F). Remarkably, in a subset of cells where the mitochondria were flattened as in mammalian cells, the ringed matrix labeling appeared encapsulated by both OMM and IMM markers (1F, bottom panels). These data indicate that the mitochondrial morphology of CCDC51-depleted cells is comparable to yeast Δ*mdm33* cells.

Previously, electron microscopy (EM) analysis of Δ*mdm33* cells revealed that the ring-like, hollow mitochondria visualized by fluorescence microscopy likely correspond to thin, elongated structures that are connected to dilated regions of mitochondria that contain relatively normal-appearing cristae invaginations (Messerschmitt et al., 2003). To examine mitochondrial ultrastructure of CCDC51-depleted cells in greater detail, we performed thin section EM analysis. As expected, control cells predominantly showed tubular mitochondria with characteristic cristae invaginations (Fig. 1G, left). In contrast, mitochondria in CCDC51-depleted cells were often swollen and enlarged with elongated cristae membranes (Fig. 1G, middle). In addition, we regularly observed mitochondria with internal ring structures, akin to Δ*mdm33* yeast cells (Fig 1G, right). As in the case of yeast cells, these ring structures contained two membrane bilayers (Fig. 1G, yellow arrows), and the cristae remained present in dilated portions of the organelle (Fig 1G, white arrows). The similarities in mitochondrial ultrastructure appearance between yeast Δ*mdm33* and U2OS CCDC51-depleted cells further indicate that the mitochondrial morphology defect is comparable in each cell type.

We next wanted to examine the dynamic behavior of the aberrant mitochondria formed in the absence of CCDC51. In the absence of Mdm33, the characteristic lariat structures can form independently of fission and fusion dynamics (in Δ*dnm1* Δ*fzo1* cells) (Klecker et al., 2015), raising the question of how these aberrant structures arise. We took advantage of the flattened cellular periphery in U2OS cells to examine the relative dynamics of lamellar mitochondria in CCDC51- depleted cells transfected with markers for the OMM (mCherry-OMP25) and the matrix (mito- HaloTag) using timelapse confocal microscopy. Notably, we could observe examples of lamellar mitochondria forming and resolving on a single mitochondrial tubule during relatively short periods (less than 20 seconds). These structures seemingly formed independent of mitochondrial fission and fusion (Fig. 1H, top panel; Movie S1). However, we also could observe examples of lamellar mitochondria that formed through apparent mitochondrial fusion events (Fig. 1H, middle panel; Movie S2). Additionally, we observed large lamellar structures that divided into multiple lamellar structures, seemingly in concert with the dynamic movements of the mitochondria (Fig. 1H, bottom panel; Movie S3). Thus, our data indicate that the altered mitochondrial morphology in the absence of CCDC51 is formed and resolved both independently of and in concert with the dynamic movements, fusion, and fission of mitochondria.

### Transient CCDC51 depletion leads to mitochondrial hyperfusion that precedes formation of lamellar mitochondria

Although the lamellar mitochondrial structures were the prevalent phenotype of cells stably depleted of CCDC51, we reasoned that this morphology could be a secondary consequence of long-term or near complete depletion of the protein. To examine the effect of acute depletion of CCDC51 in U2OS cells, we utilized RNAi to transiently knockdown the protein. Cells were transfected with either a scrambled siRNA oligonucleotide or two independent siRNAs targeted against CCDC51. We then treated cells with Mitotracker Red and assessed mitochondrial morphology by confocal microscopy. As in the case of CRISPRi cells, we could occasionally observe examples of lamellar mitochondria in CCDC51-depleted cells with both siRNAs (Fig. 2A- 2C; see Fig. 2A white arrow). However, surprisingly, the predominant phenotype we observed was mitochondrial hyperfusion compared to control cells, reminiscent of cells depleted of fission machinery such as Drp1 (Fig. 2A, 2C) (Smirnova et al., 1998). We also commonly noticed the appearance of small “nets” of mitochondria (Fig. 2A, see yellow arrow, Fig. 2C) that are similar to hyperfused mitochondrial networks found in yeast cells deficient for the Drp1 homolog Dnm1 (Bleazard et al., 1999; Sesaki and Jensen, 1999).

**Figure 2.**
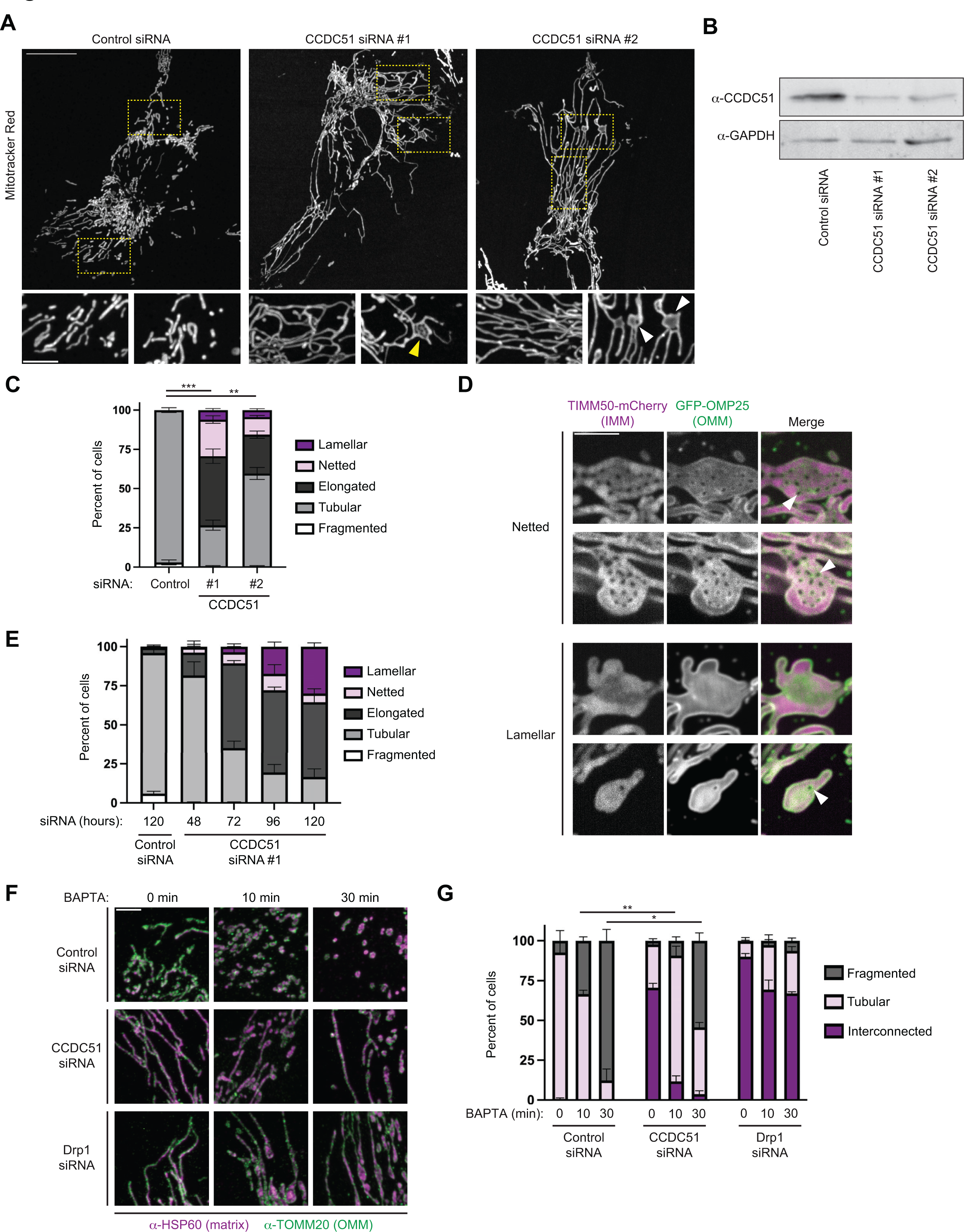
Transient CCDC51 depletion leads to mitochondrial hyperfusion that precedes formation of lamellar mitochondria. (A) Representative maximum projection confocal images are shown of U2OS cells 72h-post treatment with the indicated siRNA oligonucleotides and stained with Mitotracker Red. Insets at the bottom correspond to the indicated dashed boxes. Yellow arrows mark netted mitochondria and white arrows mark lamellar mitochondria. **(B)** Western blot analysis with the indicated antibodies of lysate from cells treated as in (A). **(C)** A graph of mitochondrial morphology characterization from the indicated siRNA-treated cells as in (A-B). Data shown represent 100 cells per condition in each of three independent experiments, and bars indicate S.E.M. Asterisks (***p<0.001; **p<0.01) represent unpaired two-tailed *t* tests of tubular mitochondrial morphology. **(D)** Representative single plane SoRa microscopy images of net-like mitochondrial morphology (top) and lamellar mitochondrial morphology (bottom) of CCDC51 siRNA-treated cells expressing TIMM50-mCherry and GFP-OMP25. Arrows mark holes that span both the OMM and IMM. **(E)** As in (C) at the indicated times post-treatment with control siRNA or CCDC51 siRNA #1. Data shown represent 50-100 cells per condition in each of three independent experiments and bars indicate S.E.M. Corresponding representative images and Western analysis are shown in Fig. S1A-S1B. **(F)** Representative maximum intensity projection confocal images are shown of cells treated with the indicated siRNA and treated with 10 µM BAPTA-AM for the indicated times. See also Fig. S1C for corresponding Western analysis. **(G)** A graph of mitochondrial morphology characterization from cells as in (F). Data shown represent 50-100 cells per condition in each of three independent experiments, and bars indicate S.E.M. Asterisks (**p<0.01; *p<0.03) represent unpaired two-tailed *t* tests of fragmented mitochondrial morphology. Scale bars: (A) 15µm (5µm in insets); (D) 5µm; (F) 4µm.

The net-like mitochondrial pattern was striking in appearance as it is a phenotype more commonly associated with defects in mitochondrial fission in yeast rather than in mammalian cells. Additionally, given the differences in staining pattern between Mitotracker and the OMM in CCDC51 CRISPRi cells (Fig. 1C vs Fig. 1E), we considered that the nets could represent an internal mitochondrial structure rather than reflect an overall net-like morphology of the organelle. Thus, to examine the membrane organization of these structures in more detail, we transiently transfected siRNA-treated cells with markers of the OMM (GFP-OMP25) and IMM (TIMM50- mCherry) and imaged by super-resolution SoRa confocal microscopy. As in the case of Mitotracker staining, we could readily visualize examples of net-like mitochondria with fluorescently tagged membrane proteins, and importantly, the “holes” of the nets were apparent with both IMM and OMM markers, indicating they span both membranes (Fig. 2D, top panels; see arrows). Conversely, lamellar mitochondria visualized with SoRa microscopy remained relatively uniformly labeled with IMM and OMM markers (Fig. 2D, bottom panels). These data indicate that the mitochondrial nets present in CCDC51 knockdown cells are akin to those in yeast Δ*dnm1* cells, and in combination with the predominant mitochondrial hyperfusion morphology, suggest that CCDC51-depleted cells have altered mitochondrial dynamics.

The presence of both lamellar and net-like mitochondrial structures in the population of CCDC51 knockdown cells led us to consider that the more severe lamellar phenotype forms due to prolonged or more complete protein depletion. To explore this possibility in more detail, we examined mitochondrial morphology in U2OS cells at 24h intervals between 48h and 120h after CCDC51 siRNA treatment. After 72h of treatment, mitochondria were predominantly hyperfused with only rare examples of mitochondrial nets and lamellar structures (Fig. 2E and Fig. S1). Continued knockdown led to an increase in the occurrence of both lamellar and net-like structures, and by 120h, mitochondria appeared lamellar relatively frequently compared to rare examples of nets (Fig. 2E and Fig. S1). We also examined the effect of CCDC51 siRNA treatment in HeLa cells to determine if the mitochondrial elongation phenotype was a general consequence of CCDC51 depletion. As in U2OS cells, mitochondria in HeLa cells became hyperfused and progressively formed lamellar structures during prolonged siRNA treatment (Fig. S2, see white arrows). Interestingly, we also commonly observed cells whose mitochondria formed enlarged bulbs (Fig. S2, see yellow arrows), a phenotype associated with mtDNA accumulation in cells defective in mitochondrial fission (Ban-Ishihara et al., 2013). Together, these data indicate that acute depletion of CCDC51 causes mitochondrial elongation and nets in the short-term, followed by the formation of lamellar structures after prolonged knockdown.

Our observations of mitochondrial hyperfusion suggest that loss of CCDC51 influences the relative frequency of mitochondrial fission and fusion dynamics. Previously, Mdm33 was proposed to aid in efficient mitochondrial division, as Δ*mdm33* cells partially resisted sodium azide-induced Dnm1-dependent fission (Klecker et al., 2015). To determine whether CCDC51- depleted cells are similarly resistant to induced mitochondrial fission, we utilized the cell permeable Ca^2+^ chelator BAPTA-AM that has been previously shown to cause Drp1-dependent mitochondrial fragmentation (Friedman et al., 2011). To minimize the influence of lamellar mitochondrial structures formed after prolonged knockdown, we treated U2OS cells with siRNA targeted against CCDC51 or Drp1 for 72 hours, which in each case led to predominantly hyperfused mitochondrial networks (Fig. 2F-2G and Fig. S1C). Cells were then mock-treated or incubated with BAPTA-AM for 10 and 30 minutes, fixed, and immunolabeled for markers of the OMM (TOMM20) and the matrix (HSP60) and imaged by confocal microscopy (Fig. 2F). In control cells, BAPTA-AM rapidly induced mitochondrial fission, and by 30 minutes, the mitochondrial network was completely fragmented in nearly all cells (Fig. 2F-2G). By contrast, in cells depleted of Drp1, mitochondrial fragmentation was nearly abolished, though mitochondrial tubules appeared shorter in a subset of cells presumably due to incomplete protein knockdown (Fig. 2F- 2G). Consistent with results in yeast Δ*mdm33* cells (Klecker et al., 2015), depletion of CCDC51 led to an intermediate phenotype but did not abolish mitochondrial fission. However, at both 10 and 30 minutes of BAPTA treatment, the proportion of CCDC51 knockdown cells with intact tubular mitochondrial networks was significantly higher than in control cells (Fig. 2F-2G). These data indicate that depletion of CCDC51 causes a delay in BAPTA-induced mitochondrial fragmentation, and together with our observations of elongated and netted mitochondria in CCDC51-depleted cells, suggest that CCDC51 is required for efficient mitochondrial fission dynamics.

### Overexpression of CCDC51 promotes Drp1-dependent mitochondrial fission

In addition to being required for normal mitochondrial morphology and dynamics, overexpression of Mdm33 causes Dnm1-dependent mitochondrial fragmentation in budding yeast (Klecker et al., 2015). We thus wanted to ask whether CCDC51 overexpression similarly causes mitochondrial fragmentation in human cells. To determine this and simultaneously examine the sub-mitochondrial distribution of CCDC51, we generated a GFP-tagged version of the protein. While CCDC51 could not be expressed when C-terminally tagged, we internally tagged CCDC51 by inserting an in-frame GFP fusion immediately after the MTS, a similar strategy that was previously used for Mdm33 (Messerschmitt et al., 2003). Importantly, this GFP-CCDC51 fusion was functional as assessed by its ability to localize to mitochondria and fully rescue the mitochondrial morphology defects of CCDC51-targeted CRISPRi cells when expressed at low levels (Fig. 3A-3B). In these cells, the GFP signal appeared distributed throughout the mitochondrial network in a patchy appearance and occasionally enriched in discrete focal structures as compared to Mitotracker staining (Fig. 3C, see arrows). To determine if the localization pattern of GFP-CCDC51 was reflective of the endogenous protein, we performed indirect immunofluorescence of wild-type U2OS cells with antibodies targeting CCDC51 and a matrix marker (HSP60) and imaged them with confocal microscopy. As in the case of GFP- CCDC51, endogenous CCDC51 was non-uniformly distributed throughout individual mitochondria and occasionally appeared concentrated at more discrete focal structures (Fig. 3D). Taken together, these data suggest that the GFP-CCDC51 construct can functionally complement the loss of CCDC51 and is reflective of the endogenous localization of CCDC51.

**Figure 3.**
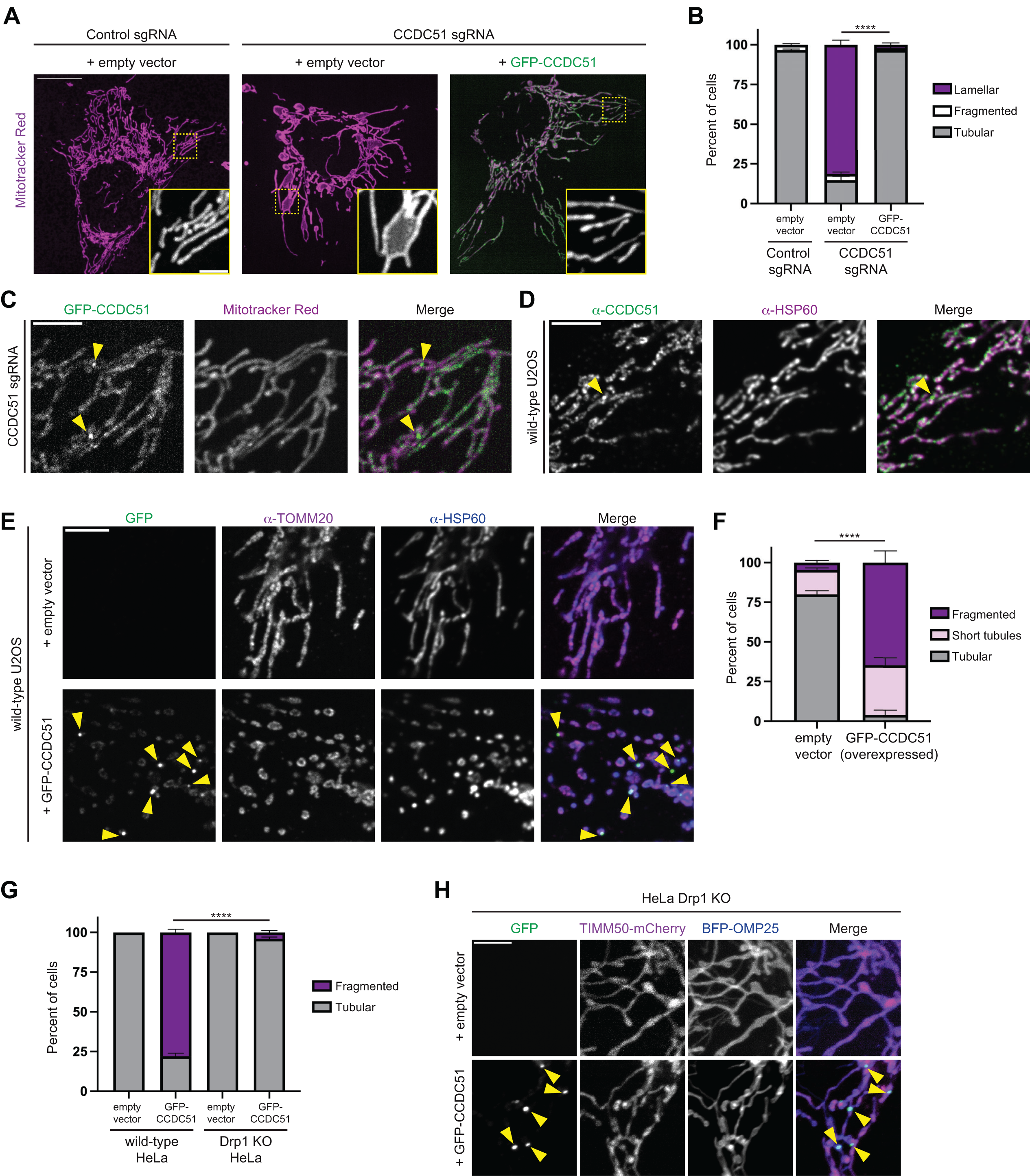
Overexpression of CCDC51 promotes Drp1-dependent mitochondrial fission. (A) Maximum projection confocal microscopy images of the indicated Mitotracker Red-stained U2OS CRISPRi cells expressing empty vector or low levels of GFP-CCDC51. Insets display Mitotracker staining corresponding to the dashed boxes. **(B)** A graph of mitochondrial morphology categorization from cells as in (A). Data shown represent 100 cells per condition in each of three independent experiments, and bars indicate S.E.M. Asterisks (***p<0.001) represent an unpaired two-tailed *t* test of tubular mitochondrial morphology. **(C)** A representative single-plane confocal microscopy image is shown of a CCDC51 CRISPRi cell, as in (A), expressing GFP-CCDC51 and labeled with Mitotracker Red. Arrows mark sites of GFP-CCDC51 concentration at discrete foci along the mitochondria. **(D)** A representative single plane confocal microscopy image is shown of a U2OS cell fixed and immunolabeled for CCDC51 and HSP60. Arrow marks a site of CCDC51 focal enrichment. **(E)** Representative single plane confocal images are shown of U2OS cells expressing empty vector (top) or high levels of GFP-CCDC51 (bottom) that were fixed and immunolabeled for TOMM20 and HSP60. Arrows mark sites of GFP-CCDC51 enrichment in discrete foci. **(F)** A graph is shown of mitochondrial morphology characterization from cells as in (E). Data shown represents 50 cells per condition in each of three independent experiments, and bars indicate S.E.M. Asterisks (****p<0.0001) represent an unpaired two-tailed *t* test of tubular mitochondrial morphology. **(G)** As in (F) for wild-type and Drp1 KO HeLa cells. Corresponding images are shown in Fig. S3A. **(H)** Representative maximum intensity confocal images are shown of HeLa Drp1 KO cells expressing empty vector (top) or high levels of GFP-CCDC51 (bottom) and co-expressing TIMM50-mCherry and BFP-OMP25. Arrows mark sites of focal accumulation of GFP-CCDC51. See also Fig. S3B for corresponding wild-type cells. Scale bars: (A) 15 µm (5µm in inset); (C) 5µm; (D) 5 µm; (E) 5 µm; (H) 5 µm.

With this CCDC51 expression construct in hand, we then assessed the consequence of its overexpression on the mitochondrial network. Wild-type U2OS cells were transfected with higher amounts of GFP-CCDC51 or an empty vector, fixed, and immunolabeled with TOMM20 and HSP60 to assess mitochondrial morphology. In cells expressing the empty vector, mitochondria in most cells were tubular in nature and the mitochondrial networks were rarely fragmented (Fig. 3E-3F). In contrast, in cells expressing high levels of GFP-CCDC51 (as established by an arbitrary fluorescence threshold), the mitochondrial morphology was drastically altered with approximately 65% of cells exhibiting fragmented mitochondrial networks (Fig. 3E- 3F), consistent with published observations (Paggio et al., 2019). Interestingly, we also observed that higher levels of GFP-CCDC51 expression correlated with an increase in its localization to discrete focal structures within the mitochondrial network (Fig. 3E, see arrows).

Next, to determine whether overexpressed GFP-CCDC51 induced mitochondrial fragmentation in a Drp1-dependent manner, we utilized HeLa Drp1 KO cells (Oshima et al., 2021). As in U2OS cells, GFP-CCDC51 expression caused mitochondrial fragmentation in wild-type HeLa cells (Fig. 3G and Fig. S3A). However, in the absence of Drp1, the mitochondrial network remained intact in nearly all GFP-CCDC51 overexpressing cells (Fig. 3G and Fig. S3A). Notably, overexpressed GFP-CCDC51 accumulated in discrete focal structures even in the absence of Drp1 (Fig. S3A, see arrows). Given this localization pattern, we also considered whether CCDC51 overexpression could lead to IMM scission independently of the OMM. To assess this, we co-transfected wild-type and Drp1 KO cells with GFP-CCDC51 as well as markers for the IMM (TIMM50-mCherry) and OMM (BFP-OMP25) (Fig. 3H and Fig. S3B). However, we found that the IMM remained continuous at the resolution of confocal microscopy, even at regions where GFP- CCDC51 enriched at foci (Fig. 3H, see arrows). Altogether, our data indicate that CCDC51 overexpression causes mitochondrial fragmentation in a Drp1-dependent manner, though CCDC51 is likely incapable of severing the IMM independently of the OMM.

### CCDC51 and Mdm33 are functional orthologs that can be spatially linked to a subset of mitochondrial division events

We have observed that acute CCDC51 depletion leads to hyperfused and net-like mitochondria, while its overexpression leads to Drp1-dependent mitochondrial fragmentation. Since both endogenous and GFP-labeled CCDC51 concentrate at discrete sites within mitochondria and the prevalence of these enrichments correlates with increased fission activity, we hypothesized that CCDC51 may be spatially linked to mitochondrial division events. To test this, we performed live cell confocal microscopy of wild-type U2OS cells co-transfected with GFP- CCDC51 and a mitochondrial matrix marker (mito-mCherry). We identified cells with low levels of GFP-CCDC51 expression and whose mitochondrial morphology remained tubular and imaged for two minutes at approximately 1 second intervals. We then identified all unambiguous mitochondrial fission events and asked whether each was spatially or temporally associated with GFP-CCDC51 localization. We analyzed a total of 78 fission events (from 18 cells) and found that 28 sites (36%) were marked by GFP-CCDC51 foci at the time of division (Fig. 4A, arrows; Movies S4-S6). Importantly, the frequency of CCDC51 localization to a division site was significantly higher than expected due to chance based on the relatively sparse distribution of GFP-CCDC51 to discrete foci along the dividing mitochondria (∼6% of the length of dividing mitochondrial tubules were covered by a foci). Strikingly, in well over half of CCDC51-marked fission events (18 of 28; 64%), the focal GFP signal dispersed simultaneously with the division event (Fig. 4A, top row). Thus, GFP-CCDC51 is both spatially and temporally associated with a subset of mitochondrial fission events in human cells.

**Figure 4.**
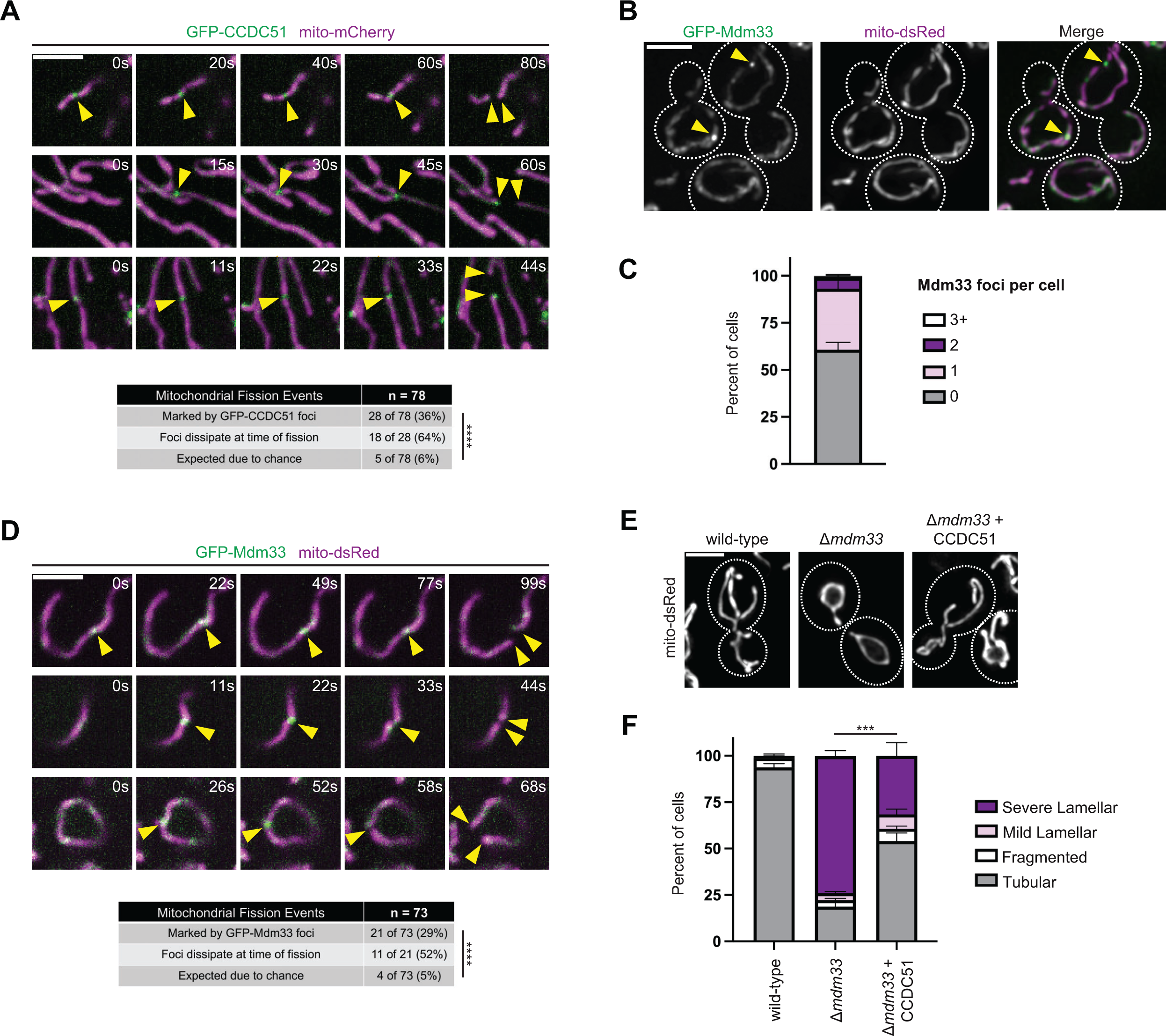
CCDC51 and Mdm33 are functional orthologs that can be spatially linked to a subset of mitochondrial division events. (A) Single-plane confocal images are shown at the indicated time intervals (s = seconds) of U2OS cells expressing low levels of GFP-CCDC51 and mito-mCherry. Arrows mark sites of GFP-CCDC51 focal enrichment relative to mitochondrial fission events. See also Movies S4-S6. The table at bottom characterizes mitochondrial fission events marked by CCDC51 and their expected frequency due to chance. Asterisks (****p<0.0001) represent a Fisher’s exact test. **(B)** A maximum intensity projection of a deconvolved fluorescence microscopy image is shown of wild-type yeast cells expressing GFP-Mdm33 from the endogenous chromosomal locus and co-expressing mito-dsRed. Dashed lines indicate cell outlines. Yellow arrows mark sites of GFP-Mdm33 focal accumulation. **(C)** A quantification of the indicated number of GFP-Mdm33 focal enrichments detected per cell as in (B). Data shown represent 75 cells per condition in each of three independent experiments and bars indicate S.E.M. **(D)** Single plane timelapse images are shown as in (A) for yeast cells expressing GFP-Mdm33 and mito-dsRed. Asterisks (****p<0.0001) represent a Fisher’s exact test. See also Movies S7-S9. **(E)** Maximum intensity projections of deconvolved fluorescence microscopy images are shown of the indicated yeast strains. Human CCDC51 is expressed under the control of an estradiol-driven promoter and all strains were grown constitutively in the presence of 20 nM estradiol. Dashed lines indicate cell outlines. **(F)** A graph of the categorization of mitochondrial morphology from cells as in (E). Data shown represent 75 cells per condition in each of three independent experiments and bars indicate S.E.M. Asterisks (***p<0.001) represent an unpaired two-tailed *t* test of tubular mitochondrial morphology. Scale bars: 3 µm.

While Mdm33 has been suggested to be involved in mitochondrial fission dynamics (Klecker et al., 2015), its localization within the mitochondrial network has not been assessed. Given the similarities between Mdm33 and CCDC51, we wanted to determine if Mdm33 has a comparable sub-mitochondrial localization pattern to CCDC51 and if it also could be spatially linked to mitochondrial fission events. To determine the localization of Mdm33, we utilized a yeast strain previously established to express functional GFP-Mdm33 from the endogenous chromosomal locus (Klecker et al., 2015). We co-expressed a matrix marker (mito-dsRed) and imaged cells by fluorescence microscopy. As in the case of GFP-CCDC51 (Fig. 3C-3D), GFP- Mdm33 displayed a non-uniform patchy distribution throughout the mitochondrial network, and occasionally enriched in discrete foci in almost half of cells at steady-state (Fig. 4B-4C, see arrows).

Next, we performed live-cell microscopy to examine the dynamics of GFP-Mdm33 foci relative to mitochondrial fission events. We imaged yeast cells for two minutes at approximately 5 second intervals, and again identified unambiguous mitochondrial fission events and determined if they were marked by focal enrichments of GFP-Mdm33. Remarkably, as in the case of CCDC51, nearly one-third of fission events were marked by a GFP-Mdm33 focal structure (21 of 73; 29%), a value that is significantly higher than would be predicted due to chance (∼5%) (Fig. 4D and Movies S7-S9). However, it should be noted that these numbers likely underreport the prevalence of Mdm33 enrichment at fission events due to weak GFP-Mdm33 signal and the difficulty visualizing mitochondria in yeast in a single plane of view. Additionally, GFP-Mdm33 was also temporally linked to fission events, as the focal enrichment dissipated at the time of fission in about half of the events we captured (11 of 21 events; 52%) (Fig. 4D). These data indicate that, like CCDC51, yeast Mdm33 is spatially and temporally linked to a subset of mitochondrial fission events.

Our data indicate that Mdm33 and CCDC51 share a similar localization pattern, localize to fission sites, and that their depletion and overexpression similarly impact mitochondrial morphology. To determine whether the two proteins are functionally orthologous, we examined whether human CCDC51 could complement the mitochondrial morphology defects of Δ*mdm33* yeast cells. We exogenously expressed untagged CCDC51 in Δ*mdm33* cells co-expressing mito- dsRed. The mitochondrial morphology appeared tubular in nearly all wild-type cells, while the mitochondrial matrix had a characteristic lariat appearance in approximately 75% of Δ*mdm33* cells (Fig. 4E-4F). Remarkably, in CCDC51-expressing yeast cells, the mitochondrial morphology defects were significantly alleviated, and over half of cells exhibited a tubular mitochondrial morphology (Fig. 4E-4F). Thus, our data indicate that CCDC51 can help maintain mitochondrial morphology in the absence of Mdm33 in a cross-species expression system and suggest that the proteins play functionally orthologous roles.

## Discussion

While mitochondrial internal organization is necessarily coupled to the overall shape and dynamics of the organelle, the mechanisms by which this occurs have remained elusive. Mdm33 is an IMM protein that is known to be critical to establish normal mitochondrial morphology in yeast (Messerschmitt et al., 2003). The protein has been implicated in mitochondrial fission dynamics; however, its precise role is unclear, and it was not previously thought to be conserved in metazoans (Klecker et al., 2015). Using a bioinformatic approach, we identified a distant metazoan ortholog, the IMM protein CCDC51. CCDC51 has similar topology and domain architecture to Mdm33 despite having minimal primary sequence homology. However, we find that loss or overexpression of CCDC51 phenocopies yeast cells with respective alterations in Mdm33 expression, and human CCDC51 can partially complement yeast Δ*mdm33* cells, indicating that they are bona fide orthologs.

Because of the unique elongated and lariat mitochondrial phenotype associated with loss of Mdm33 in yeast, its physical interactions, and its genetic interaction profile, Mdm33 has been proposed to couple mitochondrial homeostatic pathways to influence mitochondrial fission (Klecker et al., 2015). In support of this model, we observed that in human cells, acute depletion of CCDC51 leads to mitochondrial hyperfusion, and strikingly, that mitochondria form net-like structures that precede formation of the yeast-like lamellar phenotype. However, as in yeast, CCDC51 is not absolutely required for mitochondrial fission, though its loss delays the consequence of pharmacologically-induced mitochondrial fragmentation. Based on the mitochondrial dynamics phenotypes associated with the loss of Mdm33 and CCDC51, we examined the spatial distribution of each protein. Interestingly, Mdm33 and CCDC51 are non-uniformly distributed along the IMM and enrich in discrete foci in yeast and human cells, respectively. Using time-lapse confocal microscopy, we found that the proteins are spatially linked to a subset of mitochondrial fission events. Mdm33 and CCDC51 are also temporally associated with fission as the focally enriched protein commonly dissipated synchronously with the completion of fission. These data further support the possibility that Mdm33 and CCDC51 play a role in at least at a portion of mitochondrial fission events.

What could be the functional role of Mdm33 and CCDC51 in relation to mitochondrial dynamics? CCDC51 is proposed to act as a co-K^+^ channel along with the IMM protein ABCB8. One possibility is that CCDC51 and Mdm33 coordinate ion homeostasis to influence mitochondrial fission. However, CCDC51 is unusual in that its transmembrane domain does not contain primary sequence encoding a K^+^ selectively filter common to all other K^+^ channels (Mironenko et al., 2021; Paggio et al., 2019), and likewise, Mdm33 does not contain such a selectively filter sequence. Additionally, previous proteomic and genetic interaction analysis of yeast Mdm33 did not identify ABCB8 homologs (Klecker et al., 2015). Thus, this behavior of the protein may not be conserved or explain the mitochondrial morphology defects associated with its loss. Another possible explanation comes from genetic interaction analysis of yeast Mdm33, which shows combined alleviatory or synthetic sickness when Mdm33 and yeast lipid homeostasis pathways are altered. This led Westermann and colleagues to propose a lipid homeostatic role of Mdm33 in mitochondrial dynamics (Klecker et al., 2015). ER tubules pre-mark mitochondrial fission sites and while the ER may play multiple roles, recent work from Voeltz and colleagues suggests that in human cells, an ER phospholipid hydrolase acts on the mitochondria to influence mitochondrial dynamics (Nguyen and Voeltz, 2022). Thus, Mdm33 and CCDC51 may help coordinate lipid homeostasis with mitochondrial dynamics in a conserved manner.

A clue to the functional role of Mdm33/CCDC51 may also come from our observation that both proteins are only localized to a subset of mitochondrial fission events and that division can occur independently of Mdm33 and CCDC51. While mtDNA replication is associated with mitochondrial fission events, only a subset of fissions (approximately half) result in the nucleoid marker TFAM at both tips of nascent mitochondria (Lewis et al., 2016). Given Mdm33 and CCDC51 have extensive matrix-facing coiled-coil domains, the protein is potentially positioned to help spatially promote this coordination. Finally, it also is possible that Mdm33 and CCDC51 perform multiple functions, leading to the distinct morphology phenotypes observed as compared to other mutants of mitochondrial fission, as well as the disparate effects between acute and long-term depletion of CCDC51 in human cells. Regardless of the exact role of Mdm33 and CCDC51, the proteins are crucial for normal mitochondrial homeostasis, and the identification of an Mdm33 ortholog in humans demonstrates a need for future mechanistic and functional insights into the specific role(s) of the proteins.

## Materials and Methods

### Cell culture

U2OS and HEK293T cells (kindly provided by Jodi Nunnari) and HeLa wild-type and Drp1 KO cells (kindly provided by Mariusz Karbowski; (Oshima et al., 2021)) were cultured in DMEM (Sigma-Aldrich; D5796) that was supplemented with 10% fetal bovine serum (Sigma-Aldrich; F0926), 25mM HEPES, and 1% penicillin/streptomycin (Sigma-Aldrich; P4333). All experiments using CRISPRi cells were performed on early passages (<10) after sorting. Cell lines were routinely tested for mycoplasma contamination.

### Plasmid construction and siRNA oligonucleotides

CCDC51 sgRNA plasmids were made by annealing the following oligonucleotide pairs into pU6- sgRNA-Ef1a-Puro-T2A-BFP linearized at the BstXI/BlpI sites (Horlbeck et al., 2016):

CCDC51 sgRNA #1 fwd: 5’-TTGGGCACTGCAGGTAGACAGCAGTTTAAGAGC-3’ CCDC51 sgRNA #1 rev: 5’-TTAGCTCTTAAACTGCTGTCTACCTGCAGTGCCCAACAAG-3’ CCDC51 sgRNA #2 fwd: 5’-TTGGTCGGGCCACGCCAGGTACGGTTTAAGAGC-3’ CCDC51 sgRNA #2 rev: 5’-TTAGCTCTTAAACCGTACCTGGCGTGGCCCGACCAACAAG-3’ GFP-CCDC51 was generated by cloning the MTS of CCDC51, AcGFP amplified from pAcGFP1- C1, and the CCDC51 ORF from human cDNA into the XhoI/NotI sites of pAcGFP1-N1 by Gibson Assembly. GFP-OMP25 was a gift of Gia Voeltz (Addgene 141150). BFP-OMP25 and mCherry- OMP25 were generated by cloning the OMP25 cassette from GFP-OMP25 into the XhoI/BamHI sites of pTagBFP-C and pmCherry-C1, respectively. TIMM50-GFP was generated by PCR amplifying the TIMM50 cassette from pLenti-mTIMM50-mRFP (a gift of Adam Hughes; (Schuler et al., 2020)) and cloning into the XhoI/BamHI sites of pAcGFP1-N1. TIMM50-mCherry was subsequently generated by digesting the mCherry cassette from pmCherry-N1 and cloning into the BamHI/NotI sites of TIMM50-GFP. To visualize the mitochondrial matrix, Halo-MTS (referred to as mito-HaloTag; a gift of Jin Wang; Addgene 124315) and mito-mCherry (Kumar et al., 2023) were used. pcDNA3 was used as the empty vector construct in all experiments.

For transient knockdowns, the following siRNAs were used: Negative control no. 2 (ThermoFisher Scientific; 4390846), CCDC51 oligo #1 (Thermo Fisher Scientific; s36162), CCDC51 oligo #2 (Thermo Fisher Scientific; s36164), and DNM1L/Drp1 (Thermo Fisher Scientific; s19560). CCDC51 oligo #1 was used in all experiments unless stated otherwise. Specific targeted sequences are as follows: CCDC51 #1 s36162: 5’-AGACTTGGTGGGACAGATATT-3’ CCDC51 #2 s36164: 5’-GACTCAACGAGGTTCGAGATT-3’ Drp1 s19560: 5’-GACTTGTCTTCTTCGTAAATT-3’

### U2OS CRISPRi cell generation

Stable CCCD51 CRISPR interference cells were generated as previously described (Le Vasseur et al., 2021). Briefly, HEK293T cells were transiently transfected with standard lentiviral packaging plasmids and the CCDC51 sgRNA plasmids described above. Viral supernatant was harvested and filtered through a 0.45 µm PES filter and added to U2OS dCas9 cells with 5.33ug/mL polybrene. After infection and recovery, the top 50% brightest TagBFP-expressing cells were sorted by FACS. U2OS CRISPRi control sgRNA-expressing cells were described previously (Le Vasseur et al., 2021). CCDC51 sgRNA #2 was used in all experiments unless stated otherwise.

### Transient transfections

Approximately 200,000 cells were seeded per well of a 6-well dish and incubated overnight prior to transfection. Plasmid transfections were carried out with Lipofectamine 3000 (ThermoFisher Scientific) according to manufacturer’s instructions for 5-6 hours. For standard siRNA treatments, transfections were performed in 6-well plates with Lipofectamine RNAiMAX (ThermoFisher Scientific) and 20 nM RNAi oligonucleotides according to manufactuer’s instructions. Cells were incubated for ∼24 hours, passaged 1:2, grown an additional 24 hours, and transfected again for 5 hours with Lipofectamine 3000 with 20 nM RNAi oligonucleotides and any additional plasmids, where indicated. Cells were then passaged into glass-bottom microscope dishes or standard growth dishes and incubated for ∼24 hours prior to imaging or harvest for Western analysis. For knockdown time-course experiments, cells were treated as above and were imaged or harvested for lysate generation after a single transfection (48h, 72h) or after two rounds of transfection (96h, 120h).

### Mitochondrial labeling and drug treatments

To visualize mitochondria with Mitotracker, cells were stained with 25nM MitoTracker CMXRos (Thermo Fisher Scientific; M7512) in DMEM for 30-60 minutes at 37°C, washed 2x, and imaged live. To label cells transfected with mito-HaloTag, cells were stained with 1 µM Janelia Fluor 646 (Promega) for 30 minutes at 37°C and washed 2x in growth media prior to imaging. For BAPTA-AM treatment, cells were incubated with 10 µM BAPTA-AM (Calbiochem; 196419) in DMEM for 10 or 30 minutes prior to processing for immunofluorescence as described below.

### Immunofluorescence

Cells cultured on glass-bottom dishes were fixed in 4% paraformaldehyde in PBS for 15 minutes at room temperature. Cells were permeabilized for 5 minutes with 0.1% Triton X-100 in PBS, rinsed with PBS, and incubated in blocking solution for 30 minutes (PBS supplemented with 0.1% Triton X-100 and 10% FBS). Cells were then incubated with the following primary antibodies, where indicated, in blocking solution for 30-60 minutes: rabbit anti-CCDC51 (Proteintech; 20465- 1-AP), mouse anti-TOMM20 (Abcam; ab56783), mouse anti-HSP60 (Proteintech; 66041-1-Ig), and rabbit anti-HSP60 (Proteintech; 15282-1-AP). Cells were washed several times with PBS and subsequently incubated with secondary antibody with the following antibodies in blocking solution for 30-60 minutes: donkey anti-rabbit Alexa Fluor 488 (Themo Fisher Scientific; A-21206), donkey anti-mouse Alexa Fluor 555 (Thermo Fisher Scientific; A-31570), donkey anti-mouse Alexa Fluor 647 (Thermo Fisher Scientific; A-31571), or donkey anti-rabbit Alexa Fluor 647 Plus (Thermo Fisher Scientific; A32795). Cells were washed several times with PBS prior to imaging.

### Whole-cell lysate preparation and western blots

Trypsinized cell pellets were washed and lysed with freshly prepared 1x RIPA buffer (50 mM Tris-HCl pH 7.5, 150 mM NaCl, 1% Na-deoxycholate, 0.1% SDS, 1% NP-40, 1 mM EDTA) containing 1x protease inhibitor cocktail (Sigma-Aldrich; 539131) and incubated on ice for 30 minutes. Lysates were subjected to centrifugation (13,000g, 4°C, 10 minutes), supernatant was collected, and protein concentration was determined by a Bradford or BCA assay and normalized between samples. 6x Laemmli sample buffer (6% SDS, 21.6% glycerol, 0.18 M pH 6 Tris-HCl, and bromophenol blue) supplemented with 10% β-mercaptoethanol was added to a final concentration of 1x. Samples were incubated at 95°C for 5 minutes and lysates were resolved by electrophoresis on Tris-Glycine polyacrylamide gels. Protein was transferred to 0.45µm PVDF membranes and immunoblotted with the following primary antibodies, where indicated: rabbit anti- CCDC51 (1:2500; Proteintech; 20465-1-AP), mouse anti-GAPDH (1:2500; Proteintech; 60004-1- Ig), or mouse anti-DLP1 (1:1000; BD Biosciences 611112). The following secondary antibodies were used for detection: goat anti-rabbit DyLight 800 (1:10000; Thermo Fisher Scientific; SA5- 35571), goat anti-mouse DyLight 800 (1:10000; Thermo Fisher Scientific SA5-35521), goat anti-mouse DyLight 680 (1:10000; Thermo Fisher Scientific; SA5-35518), or anti-rabbit DyLight 680 (1:10000; Thermo Fisher Scientific 35568). Images were acquired by a LI-COR Odyssey Infrared Imaging System or a BioRad ChemiDoc Imaging System, and linear adjustments to images were performed using Adobe Photoshop.

### Yeast strain generation and cell growth

*Saccharomyces cerevisiae* strains were constructed in the W303 genetic background (*ade2-1*; *leu2-3*; *his3-11, 15*; *trp1-1*; *ura3-1*; *can1-100*). Routine cell growth was performed in YPD (1% yeast extract, 2% peptone, 2% glucose) or synthetic complete dextrose (SCD; 2% glucose, 0.7% yeast nitrogen base, amino acids). Strains expressing chromosomally tagged Pam17-EGFP and Tom20-mCherry were described previously (Connor et al., 2023). *MDM33* deletion was performed by PCR-based homologous recombination and the entire ORF was replaced with the HIS or NatMX6 cassettes from pFA6a series plasmids using lithium acetate transformation.

A strain expressing GFP-Mdm33 integrated at the endogenous locus was a kind gift of Jodi Nunnari (Klecker et al., 2015). To visualize the mitochondrial matrix in yeast cells, pRS304 mito-TagBFP, pRS304 mito-dsRed, and pYX142-DsRed were used (Connor et al., 2023; Friedman et al., 2011). To express human CCDC51 in yeast cells, pRS306 GalLprom-CCDC51-ADH1term was generated by Gibson assembly. This plasmid was linearized and integrated at the *ura3-1* locus. To control expression of CCDC51 with estradiol, pAGL (Veatch et al., 2009) was linearized and integrated at the *leu2-3* locus. CCDC51 expression was controlled by supplementing growth media with 20nM β-estradiol (Calbiochem 3301).

*S. cerevisiae* cells were grown at 30°C in SCD growth media to exponential phase (where indicated, in the presence of estradiol), concentrated, and plated on a 3% low melting agarose pad in SCD media on depression microscope slides. Cells were subsequently imaged as described below.

### Epifluorescence and confocal microscopy and analysis

All U2OS and HeLa imaging experiments were performed on cells adhered to glass-bottom dishes (CellVis D35-14-1.5-N or MatTek P35G-1.5-1-4C), and all live cell imaging was performed at 37°C. Epifluorescence microscopy (Figures 4B, 4E, and Figures S1-S2) was performed with a Nikon Eclipse Ti inverted epifluorescence microscope equipped with a Hamamatsu Orca-Fusion sCMOS camera, a Nikon 100x 1.45 NA objective, and an environmental control chamber. Z-series images were acquired with Nikon Elements software with a 0.2 um step size, and all images were further deconvolved using AutoQuantX 3.1 (10 iterations, blind deconvolution, and low noise). Confocal microscopy (all other figures) was performed with a Nikon Spinning Disk Confocal Microscope equipped with a Yokagawa CSU-W1, a Hamamatsu Orca-Fusion sCMOS camera, 100x 1.45 NA objective, and an environmental control chamber. Images were acquired with the Nikon Elements software using the standard spinning disk module, or where indicated, a super-resolution SoRa module. Confocal z-series were acquired with a 0.2 um step size, except where noted below. Acquisition settings were kept the same for non-transfected control cells or empty vector transfected cells within the same experiments. Linear adjustments to all images and all subsequent analyses were performed using ImageJ/Fiji.

#### The relationship between Mdm33 and CCDC51 foci and mitochondrial fission events

To assess GFP-Mdm33 foci dynamics, two-minute time-lapse movies were taken at approximately 5 second intervals with three z-planes centered on the midplane of yeast cells with a 0.3 or 0.4 µm step size. To assess whether Mdm33 foci localize to sites of mitochondrial division, maximum intensity projections were generated, and fission events were identified blind to the presence of GFP-Mdm33 foci. Single plane images were then evaluated manually for the presence and spatial relationship to GFP-Mdm33 focal accumulations.

To assess CCDC51 foci relative to mitochondrial fission events, two-minute single plane time-lapse movies were taken at approximately 1 second intervals. Cells were chosen for analysis only if GFP-CCDC51 expression was below an arbitrary fluorescence intensity threshold and mitochondrial morphology appeared tubular. Fission events were identified blind to the presence of GFP signal, and then were evaluated manually for the presence and spatial relationship to GFP-CCDC51 focal accumulations.

To determine the probability a GFP-CCDC51 or a GFP-Mdm33 foci would appear at a division site by chance, we measured the length of the dividing mitochondria tubule at the timepoint immediately preceding fission in a single plane of view. The length of the mitochondrial tubule that was covered by any discrete GFP foci was then computed and the probability was determined as the ratio between the two.

#### Mitochondrial morphology assessment

In all cases, sample identity was blinded before assessment, and cells were manually categorized as indicated in figures and legends. In the case of cells with mixed mitochondrial morphologies, cells were categorized with the more prominent morphology. In the case of yeast samples, multiple fields of view were acquired per experiment and no more than approximately 25 cells were assessed per field of view per experiment.

### Electron microscopy

To assess mitochondrial ultrastructure by EM, 75000 cells were plated onto MatTek glass-bottom dishes, incubated overnight, and fixed with 2.5% (v/v) glutaraldehyde in 0.1M sodium cacodylate buffer before submitting to the UTSW Electron Microscopy Core Facility for processing as described previously (Gok et al., 2023). After five rinses in 0.1 M sodium cacodylate buffer, cells were post-fixed in 1% osmium tetroxide and 0.8 % K3[Fe(CN6)] in 0.1 M sodium cacodylate buffer for 1h at 4°C. Cells were rinsed with water and en bloc stained with 2% aqueous uranyl acetate overnight at 4°C. After five rinses with water, specimens were dehydrated with increasing concentration of ethanol at 4°C, infiltrated with Embed-812 resin and polymerized in a 60°C oven overnight. Embed-812 discs were removed from MatTek plastic housing by submerging the dish in liquid nitrogen. Pieces of the disc were glued to blanks with super glue and blocks were sectioned with a diamond knife (Diatome) on a Leica Ultracut UCT (7) ultramicrotome (Leica Microsystems) and collected onto copper grids and post-stained with 2% uranyl acetate in water and lead citrate. Images were acquired on a JEM-1400 Plus transmission electron microscope equipped with a LaB6 source operated at 120 kV using an AMT-BioSprint 16M CCD camera.

### Bioinformatic analysis

To find putative metazoan homologs to Mdm33, HHPRED (Zimmermann et al., 2018) was used. An initial search of *S. cerevisiae* Mdm33 against the *H. sapiens* proteome with default settings identified CCDC51 as the top hit (E-value 2.7E-9 with next closest E-value 0.22). A repeat analysis with “global realignment” again identified CCDC51 (E-value 4.9E-10) as the top hit. An inverse query of the protein sequence of CCDC51 against the proteomes of *S. cerevisiae* and *S. pombe* with default settings also identified Mdm33/She9 (E-values 8.2E-4 and 2.3E-4, respectively). Transmembrane domain segments of Mdm33 and CCDC51 were identified by Phobius. Coiled-coil domains in Mdm33 and CCDC51 were identified by consensus of multiple coiled-coil prediction programs.

### Statistical analysis

Statistical analysis was performed as indicated in legends with GraphPad Prism 9.5.1. Comparisons of mitochondrial morphology were performed by two-tailed student’s *t* test of the indicated mitochondrial morphology category. Comparisons of the relationship of GFP-CCDC51 or GFP-Mdm33 foci to fission sites versus the predicted frequency due to chance were tested by Fisher’s exact test.

## Acknowledgments

We thank Laura Lackner (Northwestern University) and members of the Friedman lab for critical reading of the manuscript and helpful discussions. We thank Marcel Mettlen and The UT Southwestern Quantitative Light Microscopy Facility, which is supported in part by NIH P30CA142543, for technical advice and access to the Nikon SoRa microscope (purchased with NIH 1S10OD028630-01 to Kate Luby-Phelps) and deconvolution software. We thank the UT Southwestern Electron Microscopy Core Facility (supported by NIH 1S10OD021685-01A1) for sample preparation for EM. This work was supported by a grant from the NIH to JF (R35GM137894) with additional support for AE from the NIGMS Diversity Supplement Program. The authors declare no competing financial interests.

## Supplemental Data

**Figure S1.**
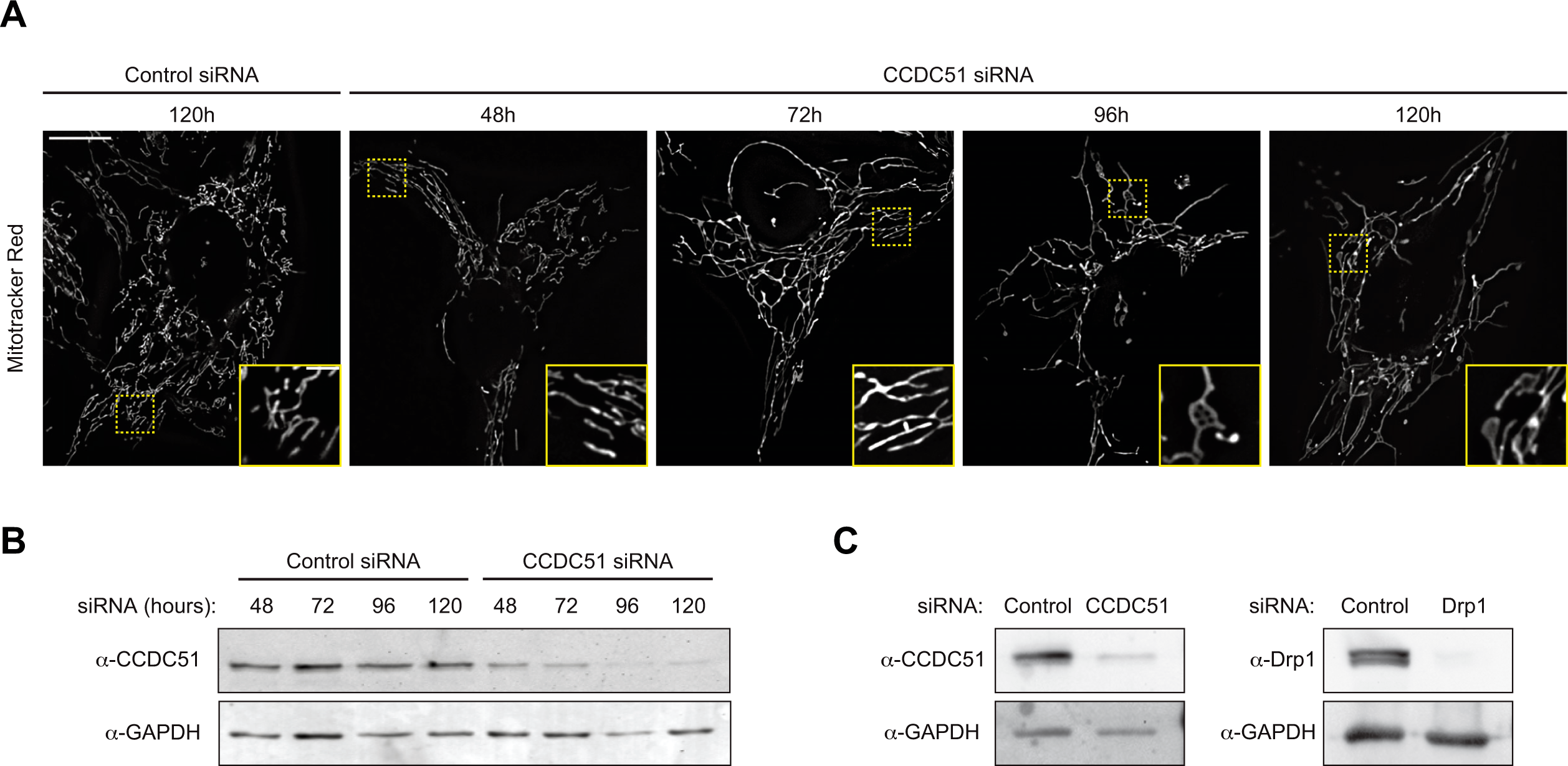
Prolonged CCDC51 depletion leads to mitochondrial hyperfusion that precedes formation of lamellar mitochondria in U2OS cells. (A) Single plane deconvolved images are shown of U2OS cells at the indicated times post-treatment with the indicated siRNA oligonucleotides and stained with Mitotracker Red. Insets correspond to dashed boxes. Images correspond to quantification shown in Fig. 2E. **(B)** Western blot analysis with the indicated antibodies of lysate from cells treated as in (A) and Fig. 2E. **(C)** Western blot analysis with the indicated antibodies of lysate from cells corresponding to Fig. 2F-2G. Scale bar: (A) 15 µm (3 µm in inset).

**Figure S2.**
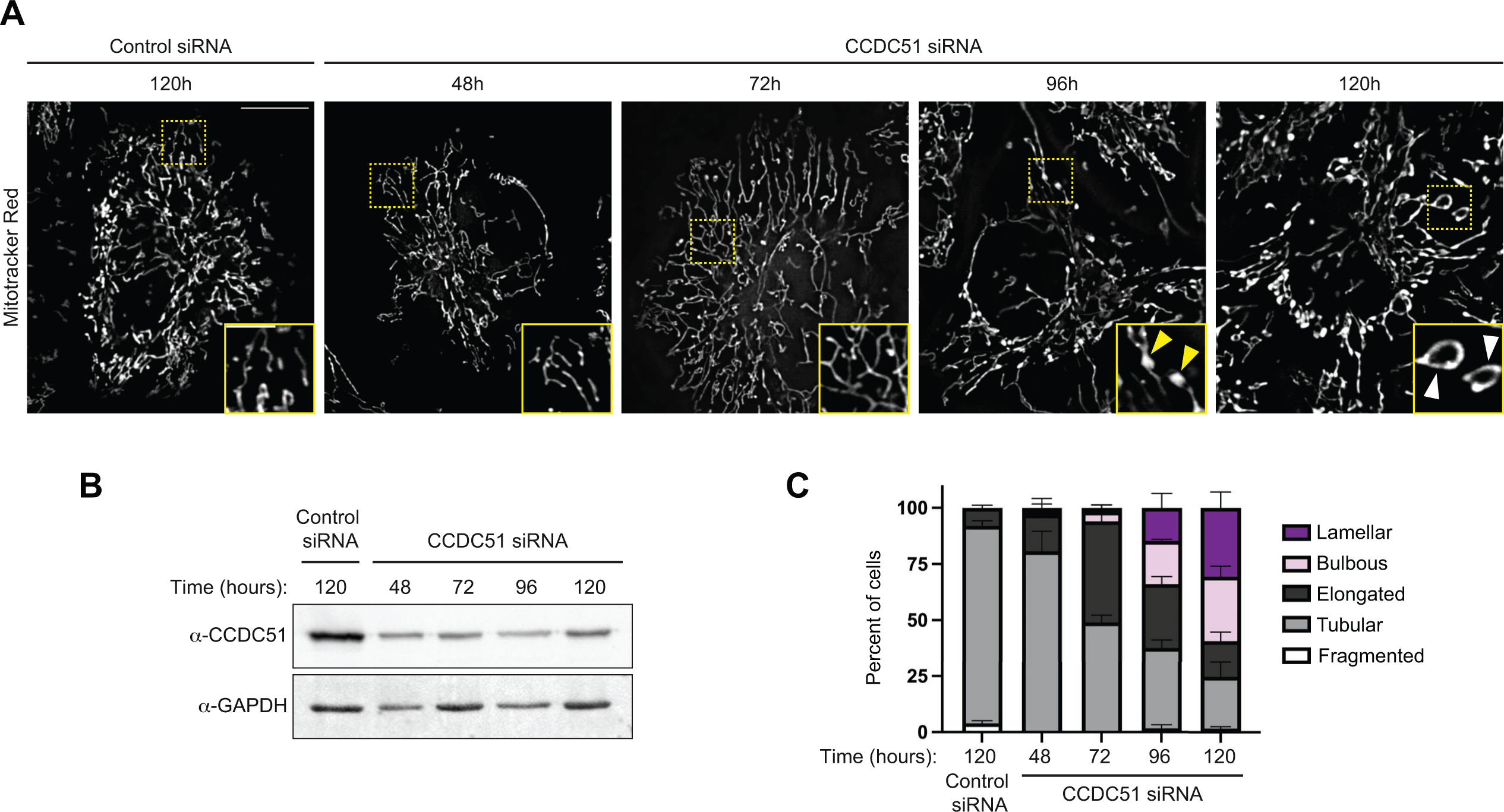
CCDC51 depletion in HeLa cells leads to mitochondrial hyperfusion and other morphological defects. (A) Single plane deconvolved images are shown of HeLa cells at the indicated times post-treatment with the indicated siRNA oligonucleotides and stained with Mitotracker Red. Insets correspond to dashed boxes. Yellow arrows mark mitochondrial bulbs and white arrows mark lamellar mitochondria. **(B)** Western blot analysis with the indicated antibodies of whole cell lysate from cells treated as in (A). **(C)** A graph showing mitochondrial morphology characterization from cells as in (A-B). Data shown represents 50-100 cells per condition in each of three independent experiments, and bars indicate S.E.M. Scale bar: (A) 15 µm (5 µm in inset).

**Figure S3.**
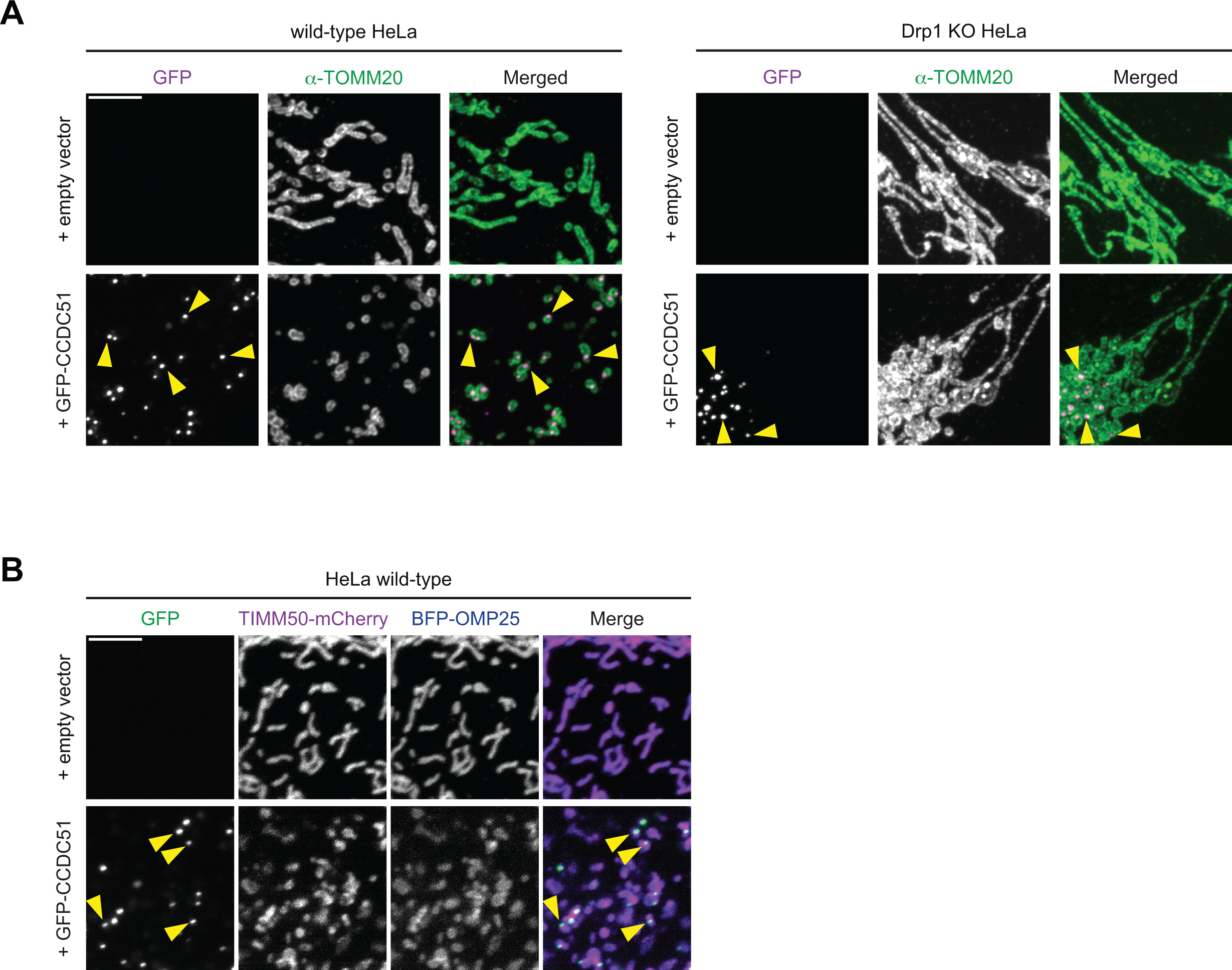
Overexpression of GFP-CCDC51 in HeLa cells leads to the fragmentation of mitochondria in a Drp1-dependent manner. (A) Representative maximum projection confocal images of HeLa wild-type and Drp1 KO cells expressing empty vector (top) or over-expressing GFP-CDCC51 (bottom) that were fixed and immunolabeled for TOMM20. Images shown correspond to quantification shown in Fig. 3G. Arrows mark sites of GFP-CCDC51 focal enrichment. **(B)** Representative maximum projection confocal images are shown of wild-type HeLa cells expressing empty vector (top) or GFP-CCDC51 (bottom) and co-expressing TIMM50- mCherry and BFP-OMP25. Arrows mark sites of focal accumulation of GFP-CCDC51. See Fig. 3H for corresponding images in HeLa Drp1 KO cells. Scale bars: (A) 5 µm; (B) 5 µm.

